# Dominant *RDH12*-retinitis pigmentosa impairs photoreceptor development and cone function in retinal organoids

**DOI:** 10.1101/2024.12.11.627489

**Authors:** Cécile Méjécase, Ya Zhou, Nicholas Owen, Pablo Soro-Barrio, Riccardo Cheloni, Neelima Nair, Hajrah Sarkar, Lyes Toualbi, Mariya Moosajee

## Abstract

Retinal dehydrogenase 12 (RDH12) is a photoreceptor NADPH-dependent retinal reductase enzyme, converting all-*trans*-retinal to all-*trans*-retinol. Heterozygous variants in *RDH12* cause a rare autosomal dominant (AD) retinitis pigmentosa. As no disease models exist, we generated human induced pluripotent stem cell derived retinal organoids (RO) from a *RDH12-*AD patient (with pathogenic c.759del p.(Phe254Leufs*24) variant), alongside a healthy control (WT). *RDH12-*AD RO exhibited correct localisation of RDH12 to the photoreceptor inner segments up to week 44; transmission electron microscopy at week 37 showed photoreceptors were less abundant and shorter in length compared to WT. Visual cone function, retinol biosynthesis and the vitamin A pathway were also highly disrupted at week 44. Our study is the first to describe a *RDH12*-AD disease model with pathology at later stages of photoreceptor differentiation, in keeping with the milder disease course seen in humans. It provides insights into the aetiology and possible targets for future therapeutic development.

## 1 Introduction

Inherited retinal diseases (IRDs) are a genetically and heterogeneous group of non-progressive and progressive sight loss disorders. Among the progressive group, the severity of the clinical phenotype can vary from a severe early onset Leber congenital amaurosis (LCA) to a mild adult-onset retinitis pigmentosa arising from variants in the same causative gene (Georgiou et al., 2021). *RDH12* is a key example, ∼80 biallelic pathogenic variants account for up to 10% of LCA (Sarkar and Moosajee, 2019), but there are 4 rare cases of autosomal dominant (AD) late-onset retinitis pigmentosa (RP) reported (Fingert et al., 2008; Muthiah et al., 2022; Sarkar et al., 2020).

*RDH12* encodes the retinal dehydrogenase 12 protein, a NADPH-dependent retinal reductase enzyme (Sarkar and Moosajee, 2019). Localised to the inner segment of photoreceptors (Kurth et al., 2007), RDH12 is involved in visual cycle. In photoreceptor outer segments, rhodopsin interacts with 11-*cis*-retinal and its photoactivation transforms it into all-*trans*-retinal. Most of all-*trans*-retinal is reduced into all-*trans*-retinol by RDH8 in photoreceptor outer segments, the remaining is processed by RDH12 in the inner segment to protect cells from oxidative stress (Sarkar and Moosajee, 2019). RDH12 has also been reported to metabolise medium-chain aldehyde, especially nonanal (Belyaeva et al., 2005), to resist to induced toxicity in HEK293 models (Lee et al., 2008).

While few *in vitro* (cell lines overexpression WT or mutated *RDH12*) and *in vivo* (mutant mouse and zebrafish) models exist for autosomal recessive *RDH12*-retinopathies, no animal model was created for autosomal dominant *RDH12*-related retinitis pigmentosa. The heterozygous *Rdh12* mouse model does not exhibit any retinal phenotype (Kurth et al., 2007; Maeda et al., 2006). Previous studies report retinal organoids derived from patients cells recapitulate the phenotype observed in patients and are a good model to understand pathophysiology associated to autosomal dominant retinal disorders (Buskin et al., 2018; Chirco et al., 2021; Gao et al., 2020; Kruczek et al., 2021; Rodrigues et al., 2022). Hence, in this study, we generated and characterized retinal organoid models differentiated from human induced pluripotent stem cells (HiPSC) derived from a patient with heterozygous pathogenic frameshift c.759del p.(Phe254Leufs^*^24) in *RDH12*, who was affected with dominant mild, late-onset RP (*RDH12*-AD) (Sarkar et al., 2021, 2020).

## 2 Material and Methods

### 2.1 Cell culture

HiPSC were obtained from a RDH12-AD patient described in (Sarkar et al., 2020) and an unrelated unaffected age- and gender-matched healthy control (WT) (Méjécase et al., 2020; Sarkar et al., 2021). Three clones for each line were expanded as previously reported (Méjécase et al., 2020; Sarkar et al., 2021). HiPSC were differentiated into retinal organoids following a feeder-free xeno-free protocol (Reichman et al., 2017). Optic cups were picked between day 21 and day 32, cultured in 6-well plates with 10 ng/mL of human FGF2 during 7 days (Reichman et al., 2017). Each optic cup was selected and cultured in 96-well plate, with the previously reported protocol to obtain laminated retinal organoids (Gonzalez-Cordero et al., 2017). Retinoic acid (0.5µM) is maintained until the latest timepoint (week 44).

### 2.2 Retinal organoid characterisation and electron microscopy analysis

Retinal organoids were cryoembedded at two timepoints (week 18 and week 44) for immunostaining as previously reported (Reichman et al., 2014). Immunofluorescence staining was performed on 12-µm thick sections. Samples were incubated in a permeabilising solution (0.5% Triton X-100, 0.1% Tween20, PBS) for 1 hour, in a blocking solution (PBS, 0.2% gelatin, 0.25% Triton X-100) for 1 h at room temperature, and then in primary antibodies solutions diluted in blocking solution (**Supplementary Table S1**) overnight at 4°C. After washes in PBS, 0.1% Tween20 solution, sections were incubated with secondary antibodies (**Supplementary Table S1**) diluted in blocking solution during 1 h at room temperature. Sections were mounted with ProLong Diamond Antifade Mountant with DAPI (ThermoFisher).

A TUNEL assay was performed, using the ApopTag Fluorescein In situ Apoptosis detection kit (Millipore) as per the manufacturer’s instructions. The entire organoid section (19-28 images per stack) was imaged using LSM710 confocal microscope at magnification x40 and a maximum intensity projection Z stack was performed for each image (Lane et al., 2020). TUNEL positive nuclei were count manually on Fiji. After a mean filter (radius = 2.00), Li threshold and Watershed binary treatment, particles with a minimum size of 3 µm and a circularity between 0.20 to 1.00 were counted as total nuclei with DAPI staining. Approximately 820 nuclei were counted with a maximum of 29 TUNEL positive cells.

Two retinal organoids per line (week 37) were fixed with Karnovsky’s fixative (2.5% Glutaraldehyde, 1% PFA, 80 mM Cacodylate buffer (pH 7.4), 20 mM NaOH) for 1 h at room temperature in the dark. After washes in 0.2M Sodium cacodylate solution, samples were postfixed in 1% aqueous osmium tetroxide for 1-2 h at room temperature. A serial dehydration with increasing ethanol concentration (50%, 70%, 90% and 100%) was performed, followed with 100% peropylene oxide bath. Samples were then embedded in 100% EPON resin. The samples were serial sectioned at the thickness of 70nm using a UC7 ultramicrotome (Leica Microsystems) and sections were picked up on Fomvar-coated 2mm slot copper grids (Gilder Grids). The sections were post-stained with 2% UA and lead citrate, and sections containing the region of interest were imaged using a 120 kV JEM-1400Flash Transmission electron microscope with a sCMOS Flash camera (JOEL UK). For each organoid per line, three regions 60 µm apart were studied: photoreceptor segments were counted and measured from outer limiting membrane. The measurement and counting was performed blind. As no significant difference was observed between the two researchers’ analysis for three blinded regions (**Supplementary Figure S1**), the rest of the analysis (measurement and counting) was performed by only one researcher. A total of 2,097 and 1,238 segments were counted in WT and *RDH12*-AD samples. Differences were tested with a linear mixed model, including segment length as the outcome variable, condition as fixed effect and organoid number and region as random effects. Linear mixed model analysis showed a significant effect of condition on segment length (𝒳^2^=16.2, p=0.001). Post-hoc analysis showed significant pairwise differences between WT and *RDH12*-AD retinal organoids (all p<0.05, ^*^).

### 2.3 RNAseq data analysis

At week 44 (day 303), 12 replicates (two organoids per replicate) from two differentiations of two *RDH12* AD clones and 8 replicates from three differentiations from one WT clone were snap frozen. RNA was isolated, quantified and cDNA generated from pooled organoids (Qiagen, UK). cDNA libraries were sequenced on a HiSeq 2500 with paired-end reads of 150 bp using Illumina Stranded Total RNA with Ribo-Zero Plus (Illumina, UK). Raw reads were quality and adapter trimmed using cutadapt (version 3.4) (Martin, 2011) before alignment. Reads were mapped and subsequent gene-level counted using RSEM 1.3.1 (Li and Dewey, 2011) and STAR 2.7.10a (Dobin et al., 2013) against the human genome GRCh38 using annotation release 95, both from Ensembl. Normalisation of raw count data and differential expression analysis was performed with the DESeq2 package (version 1.38.3) (Love et al., 2014) within the R programming environment (version 4.2.2) (R Development Core Team, 2008). The pairwise comparison was performed with the contrast function, from which genes differentially expressed (adjusted P value being less than 0.05) between different conditions were determined. Gene lists were used to look for pathways and molecular functions with over-representation analysis using Reactome (Jassal et al., 2020) and Gene Ontology (GO) (Mi et al., 2019).

### 2.4 Statistics and reproducibility

All statistical analyses were performed using R for Statistics (https://www.r-project.org/, v4.1.2) or GraphPad Prism 8 Data presented are shown as the mean (m) ± s.d. *P* ≤ 0.05 was considered statistically significant.

### 2.5 Data availability

The RNA-seq data generated by this study have been deposited in the NCBI Gene Expression Omnibus under the access code GSE271751, including unprocessed FASTQ files and associated gene count matrices.

## 3 Results

### 3.1 Retinal organoids characterisation at different time points

HiPSC derived from *RDH12*-AD patient and WT control were differentiated into retinal organoids and characterised after 18 and 44 weeks of differentiation (**Figure 1**). At week 18, both WT and *RDH12*-AD retinal organoids developed rod and cone photoreceptors and there was no significant difference in cell death (**Figure 1A**). The *RDH12*-AD rod and cone photoreceptors presented with shorter segments than WT. At week 44, there was no significant difference in cell viability but the photoreceptor segments remained shorter in appearance in the *RDH12*-AD mutant, particularly the cones (**Figure 1B**). RDH12 was detected in the photoreceptor inner segments in both WT and *RDH12*-AD retinal organoids at week 18 and week 44.

**Figure 1:**
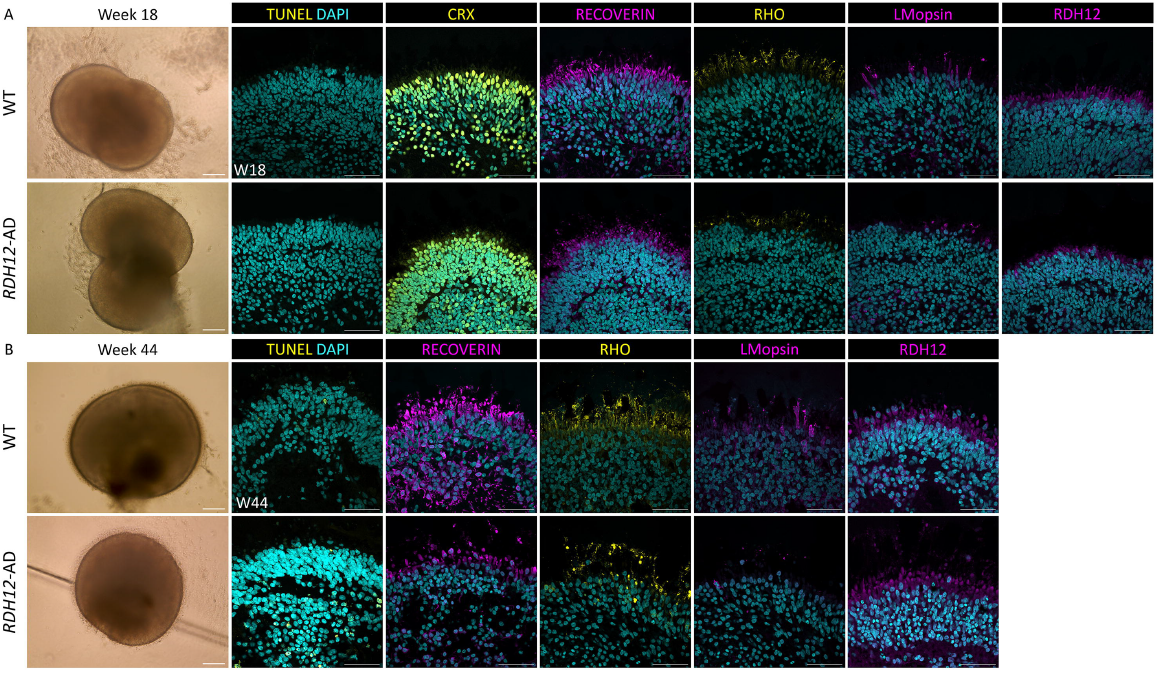
Immunofluorescence studies revealed *RDH12*-AD retinal organoids developed abnormal photoreceptors during differentiation. **(A)** *RDH12*-AD organoids developed rod and cone photoreceptors with abnormal segments compared to WT photoreceptors at week 18. N=5 per line, from 4-3 differentiation batches, from 2-3 clones. **(B)** Photoreceptors remained abnormal in mature *RDH12*-AD retinal organoids at week 44. N=3 organoids per line, from 1-2 differentiation batches, from 1-2 clones. CRX, RECOVERIN: photoreceptor precursors; RHO: rod; L/M-opsin: cone. Scale bar: 200 µm for brightfield images; 50 µm.

### 3.2 Photoreceptors are shorter and less abundant in *RDH12*-AD retinal organoids

Transmission electron microscopy of week 37 retinal organoids was undertaken to look for intracellular changes (**Figure 2**). Both WT and *RDH12*-AD organoids developed segments around the organoid edges, with mitochondria and a clear and regular outer limiting membrane, and few of them have intracellular membrane discs, similar to the outer segment structure (**Figure 2A**). Blinded analyses of three regions per organoid showed WT retinal organoids have an average 62 ± 5 photoreceptors per 100 µm whilst *RDH12*-AD have significantly less photoreceptors 33 ± 4 photoreceptors per 100 µm (p<0.05, **Figure 2B**). Moreover, average photoreceptor segment length was 10.09 ± 6.48 µm in WT (range between 0.25 µm to 44.02 µm) and reduced to 5.51 ± 3.42 µm in *RDH12*-AD retinal organoids (range between 0.36 µm to 26.77 µm) (p<0.05, **Figure 2C**).

**Figure 2:**
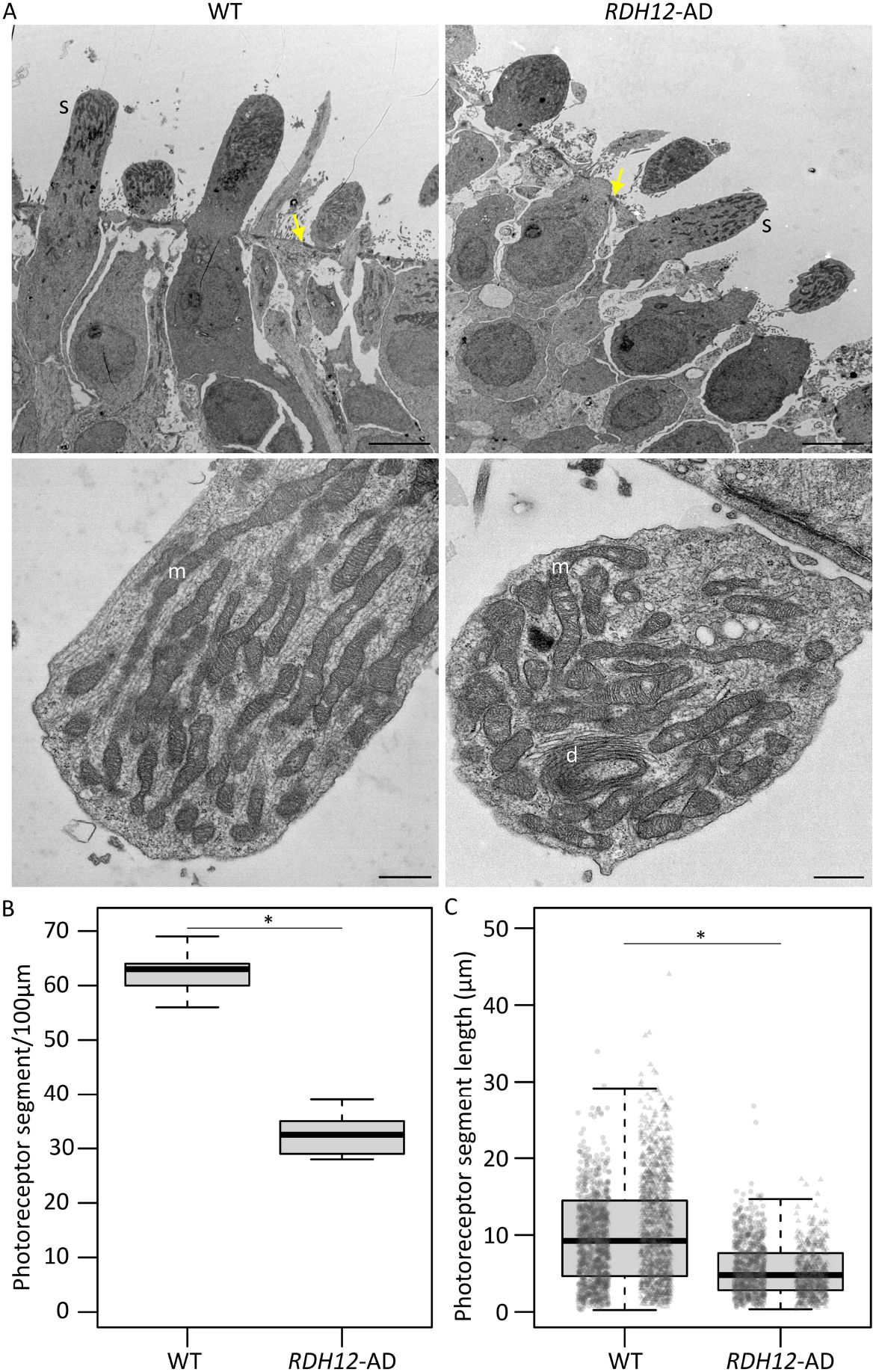
Ultrastructure of *RDH12*-AD mutants and WT retinal organoids at week 37. **(A)** Both WT and *RDH12*-AD retinal organoids developed photoreceptor segments (s), with mitochondria (m) and discs (d). Yellow arrows indicate outer limiting membrane. Scale bar: 5 µm (top), 500 nm (bottom). **(B)** *RDH12*-AD retinal organoids had significantly less photoreceptors per 100 µm than WT organoids (p<0.05). **(C)** Photoreceptor segments were significantly shorter in *RDH12*-AD retinal organoids compared to WT (p<0.05). Data are represented as mean ± SEM. N=3 regions from 2 organoids per line, from 1 differentiation batch, and a total of n=2097 and n=1238 photoreceptor segments, from outer limiting membrane to tip end, were counted and measured in WT and *RDH12*-AD organoids respectively.

### 3.3 Dominant disease-causing variant in *RDH12* impairs cone functions

Bulk RNA sequencing was performed on mature RDH12 and WT retinal organoids at week 44, to identify disease-causing pathways. In mature *RDH12*-AD retinal organoids, 2,446 genes were differentially expressed, with 1,424 upregulated and 1,022 downregulated, compared to WT expression (**Figure 3A**). Gene ontology (GO) analysis identified GO over-representation of passive transmembrane transporter activity annotations (including GO:0022803, GO:0005249, GO:0005261, GO:0046873, GO:0015103, GO:0015267, GO:0022836, GO:0005267, GO:0022843, GO:0005244, GO:0022832, GO:0015079, GO:0005216, GO:0022824, GO:0005230, GO:0022834, GO:0005231, GO:0005237, GO:0005245, GO:0005254, GO:0016917, GO:0004890, GO:0022835, GO:0015276, GO:0015085, GO:0005262, GO:0005253, GO:0015108, GO:1904315, GO:0099095, GO:0022851, GO:0008331), neurotransmitter receptor activity (including GO:0098960, GO:0099529 and GO:0008503), nucleoside-triphosphatase regulator activity (including GO:0060589, GO:0030695, GO:0005096), photoreceptor activity (including GO:0009881 and GO:0008020), lipid phosphatase activity (including GO:0042577 and GO:0008195), modified amino acid binding (GO:0072341, GO:0001786), extracellular matrix structural constituent (including GO:0005201 and GO:0030020), glycosaminoglycan binding (including GO:0005539), transmembrane receptor protein tyrosine kinase activator activity (GO:0030297), ephrin receptor activity (GO:0005003), cGMP binding (GO:0030553), SNAP receptor activity (GO:0005484), cell-cell adhesion mediator activity (GO:0098632), coreceptor activity (GO:0015026), collagen binding (GO:0005518), actin binding (GO:0003779), cadherin binding (GO:0045296), integrin binding (GO:0005178), 1-phosphatidylinositol binding (GO:0005545), tubulin binding (GO:0015631, GO:0043015) and microtubule binding (GO:0008017) in *RDH12*-AD (**Figure 3B** and **Supplementary Table S2**).

**Figure 3:**
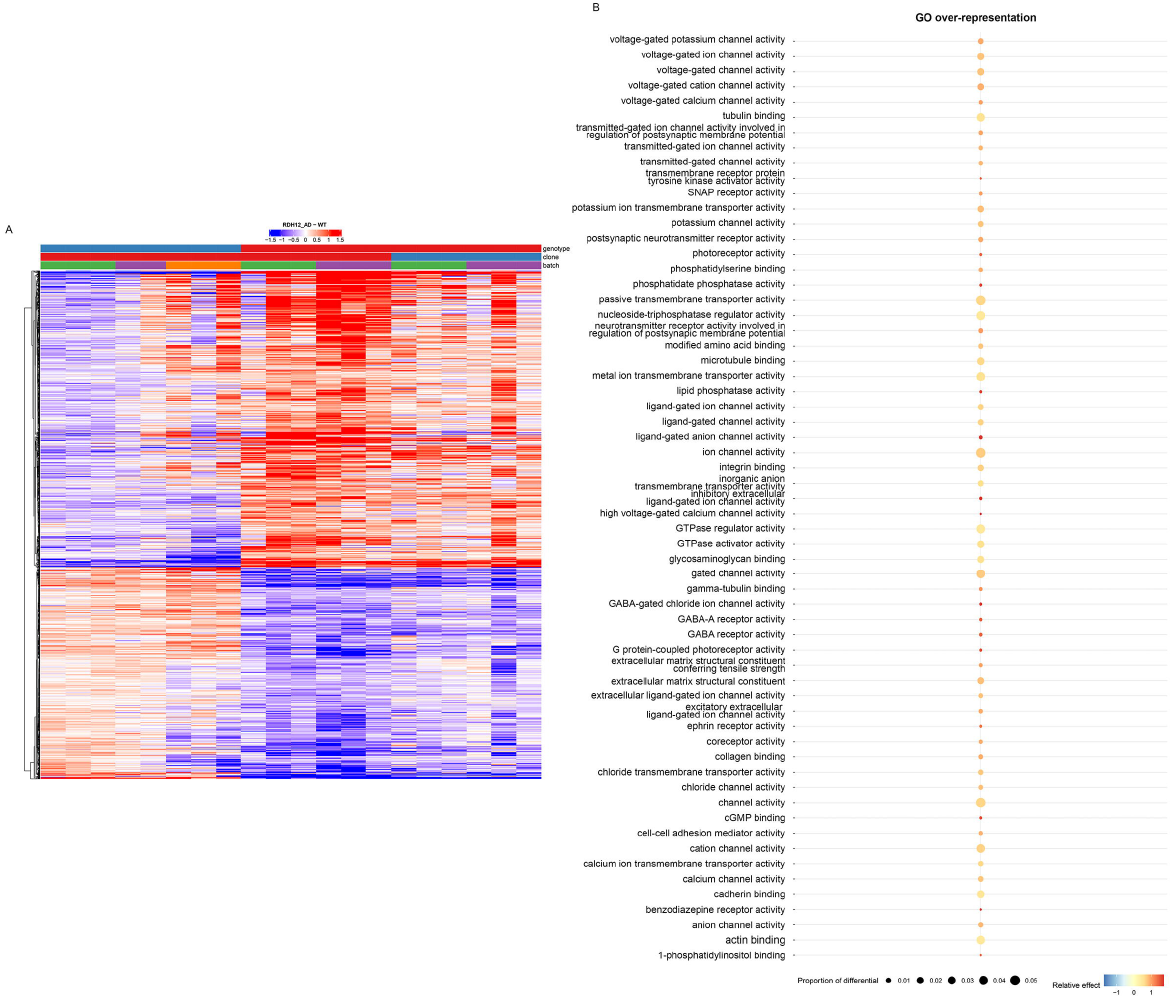
Transcriptomic analyses distinguish *RDH12*-AD mutant to WT retinal organoids and highlight molecular function affected in mutant organoids (day 303 week 44). **(A)** Heatmap of differential genes along the two samples, WT (blue) and *RDH12*-AD (red). **(B)** Gene ontology (GO) analysis identified over-represented GO molecular function in *RDH12*-AD organoids. N=12 replicates (two organoids per replicate) from two differentiations of two *RDH12* AD clones and N=8 replicates from three differentiations from one WT clone.

We examined different retinal cell markers (Kim et al., 2023), and found cone-specific markers were downregulated in *RDH12*-AD retinal organoids compared to WT: *OPN1LW, TDRG1, GRK7, RAB41, MYL4, PDE6H, CNGB3, PDE6C, GNAT2, GUCA1C, GNGT2, ARR3, SLC24A2, KCNB2* and *PEX5L* (**Supplementary Table S3 and Figure 4A**). Five ganglion cell markers (*CLEC2L, RBPMS, NEFM, SLC17A6, TMEM163*) were upregulated in *RDH12*-AD retinal organoids (**Supplementary Table S3 and Figure 4A**). However, only two stress and/or apoptosis markers were differentially expressed: *MOAP1* (LFC -0.804, padj<2.109^*^10^-8^) and *AIFM2* (LFC 0.639, padj<3.276^*^10^-2^) (Deng et al., 2018) (**Supplementary Table S3**).

**Figure 4:**
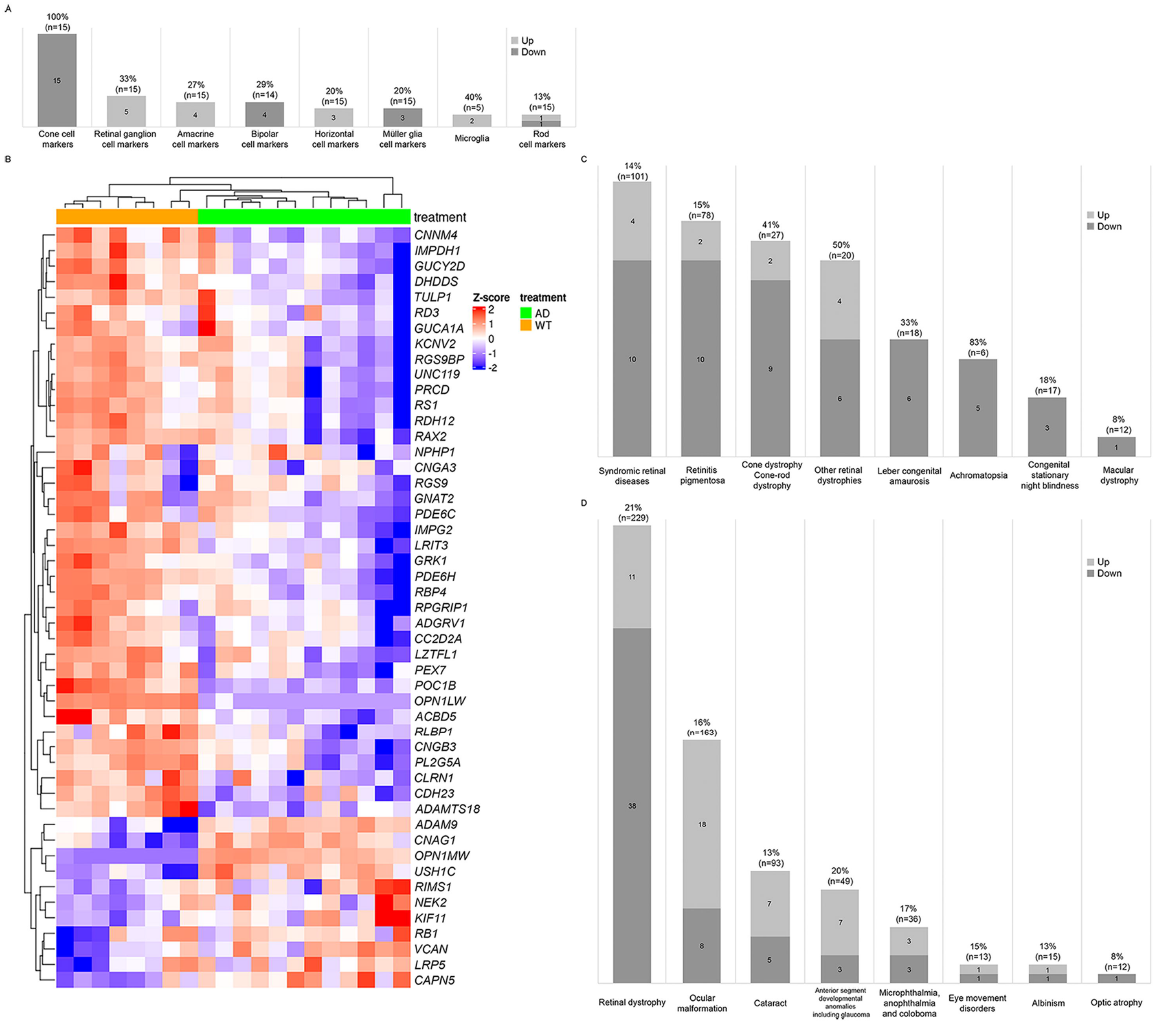
Retinal markers and genes associated with inherited eye diseases were dysregulated in *RDH12*-AD retinal organoids at day 303 (week 44). **(A)** Cone-specific cell markers were downregulated in *RDH12-*AD retinal organoids compared to WT. **(B-C)** Forty-nine unique genes associated with retinal dystrophies were significantly differentially expressed. **(D)** In total, seventy-nine unique genes associated with inherited eye diseases were significantly dysregulated in *RDH12*-AD retinal organoids. N=12 replicates (two organoids per replicate) from two differentiations of two *RDH12* AD clones and N=8 replicates from three differentiations from one WT clone.

We compared expression of genes previously reported to be associated with inherited eye diseases (**Supplementary Table S4 and Figure 4**). Interestingly, 49 genes associated with retinal dystrophies were significantly differentially expressed in *RDH12*-AD (21%, n=229), with 38 downregulated and 11 upregulated (**Figure 4B-D and Supplementary Table S4**). These genes are associated with syndromic retinal disorders (*ACBD5, ADAMTS18, ADGRV1, CAPN5, CC2D2A, CDH23, CLRN1, KIF11, LZTFL1, NPHP1, PEX7, POC1B, USH1C, VCAN*), retinitis pigmentosa (*ADAMTS18, CC2D2A, CLRN1, CNGA1, DHDDS, IMPDH1, IMPG2, NEK2, PRCD, RDH12, RLBP1, TULP1*), other retinal dystrophies (*CAPN5, LRP5, OPN1LW, OPN1MW, PLA2G5, RB1, RBP4, RGS9, RGS9BP, RS1*), Leber congenital amaurosis (*GUCY2D, IMPDH1, RD3, RDH12, RPGRIP1, TULP1*), cone dystrophy/cone-rod dystrophy (*ADAM9, CNNM4, GUCA1A, GUCY2D, KCNV2, PDE6C, POC1B, RAX2, RIMS1, RPGRIP1, UNC119*), achromatopsia (*CNGA3, CNGB3, GNAT2, PDE6C, PDE6H*), congenital stationary night blindness (*LRIT3, GRK1, RLBP1*) and macular dystrophy (*RAX2*). Differential gene expression was also observed for the different groups of ocular diseases (**Figure 4D, and Supplementary Table S4**): 26 genes associated with ocular malformation (16%, n=163), with 18 are upregulated and 8 downregulated; 12 are associated with cataract (13%, n=93), with 7 upregulated and 5 downregulated; 10 associated with anterior segment developmental anomalies including glaucoma (20%, n=49), with 7 upregulated and 3 downregulated; 6 associated with microphthalmia, anophthalmia and coloboma (17%, n=36), with 3 upregulated and 3 downregulated; 2 associated with eye movement disorders (15%, n=13), with 1 upregulated and 1 downregulated; 2 were associated with albinism (13%, n=15), with 1 upregulated and 1 downregulated; and 1 gene downregulated, associated with optic atrophy (8%, n=12).

RDH12 is involved in several pathways: vision cycle in rod and cone photoreceptors, vitamin A and carotenoid metabolism, retinol biosynthesis and metabolism (Sarkar and Moosajee, 2019). Interestingly, forty-five unique genes involved in these pathways were dysregulated in *RDH12*-AD retinal organoids compared to WT (**Figure 5A and Supplementary Table S5**): 18 genes involved in visual signal transduction in cones (78%, n=23), 16 are downregulated and 2 upregulated; 27 genes involved in visual phototransduction (25%, n=110), 17 downregulated and 10 upregulated; 22 genes involved in visual cycle in retinal rod (23%, n=94), 17 downregulated and 5 upregulated; 9 genes involved in Vitamin A and carotenoid metabolism (21%, n=42), 7 downregulated and 2 upregulated; 2 genes involved in retinol metabolism (5%, n=38) are downregulated; 4 genes involved in retinol biosynthesis are downregulated (31%, n=13). Cone function appeared to be highly affected in *RDH12*-AD retinal organoids (**Supplementary Table S5**): *GRK7* (LFC -1.974, padj<2.061^*^10^-10^), *OPN1MW3* (LFC -1.746, padj<4.244^*^10^-3^), *PDE6H* (LFC -1.541, padj<2.014^*^10^-10^), *CNGB3* (LFC - 1.514, padj<4.937^*^10^-5^), *PDE6C* (LFC -1.509, padj<1.042^*^10^-12^), *GNAT2* (LFC -1.499, padj<8.770^*^10^-9^), *GNGT2* (LFC -1.447, padj<1.042^*^10^-5^), *ARR3* (LFC -1.438, padj<1.528^*^10^-6^), *SLC24A2* (LFC -1.403, padj<1.079^*^10^-3^), *GNB3* (LFC -1.202, padj<6.145^*^10^-4^), *RDH12* (LFC - 1.158, padj<4.633^*^10^-3^), *GRK1* (LFC -1.132, padj<6.192^*^10^-5^), *RGS9BP* (LFC -0.886, padj<2.274^*^10^-3^), *CNGA3* (LFC -0.824, padj<2.127^*^10^-2^), *GUCY2D* (LFC -0.788, padj<2.388^*^10^-3^), *RGS9* (LFC -0.589, padj<3.590^*^10^-2^); and *OPN1MW* (LFC 10.42, padj<3.188^*^10^-39^), *OPN1MW2* (LFC 10.658, padj<2.172^*^10^-25^). Vitamin A pathway was downregulated in *RDH12*-AD retinal organoids (**Figure 5B-C**).

**Figure 5:**
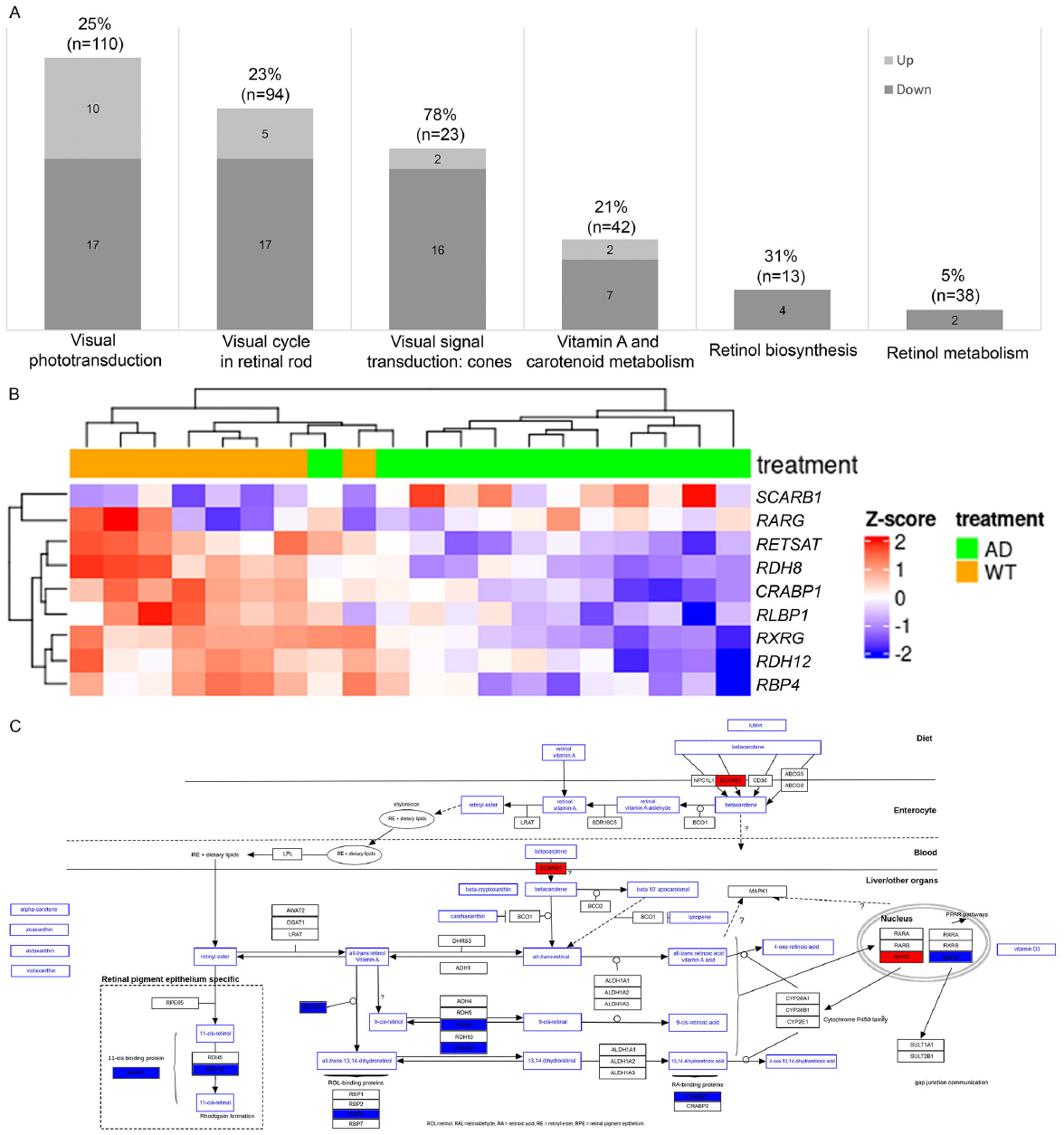
RDH12 pathways were differentially expressed in *RDH12*-AD retinal organoids at day 303 (week 44). **(A)** Forty-five genes involved in RDH12 pathways were significantly differently expressed. **(B-C)** Vitamin A pathway was affected in *RDH12*-AD retinal organoids at day 303 (W44), genes in blue are downregulated; genes in red upregulated in mutant organoids. N=12 replicates (two organoids per replicate) from two differentiations of two *RDH12* AD clones and N=8 replicates from three differentiations from one WT clone.

## 4 Discussion

This study describes the first retinal organoid model for autosomal dominant *RDH12*-related disease, derived from a patient with a heterozygous frameshift variant c.759del p.(Phe254Leufs*24). This powerful tool unravels disease-mechanisms associated with a milder and late-onset retinitis pigmentosa, providing further insights of RDH12 function in the retina.

Autosomal dominant *RDH12* retinopathy is considered a milder form of IRD with relatively late-onset in adulthood. It is characterised by nyctalopia and progressive visual field loss with associated bone spicules in peripheral retina and retinal vessel attenuation, central vision is affected much later in the disease, although a number of peripheral cone photoreceptors are reduced and their outer segment is abnormal (Fingert et al., 2008; Muthiah et al., 2022; Sarkar et al., 2020). *RDH12*-AD retinal organoids model the mild phenotype seen in patients; they display a similar development to WT organoids with production of the outer nuclear layer with mature photoreceptors and their segments. Photoreceptor development is normal in *RDH12*-AD retinal organoids; the expression of photoreceptor progenitor and photoreceptor transcription markers (*CRX, OTX2, PAX6, NRL, LHX2, NR2E3, NEUROD1, VSX2, PRDM1, ROM1* (Zhao and Peng, 2021)) remain unchanged.

Interestingly, *LRIT3, GRK1* and *RLBP1*, previously reported to be associated with autosomal recessive or X-linked congenital stationary night blindness (Zeitz et al., 2015), are downregulated in the *RDH12*-AD retinal organoids, which explain the presenting nyctalopia symptoms observed in the two unrelated families with autosomal dominant *RDH12*-RP (Sarkar et al., 2020). Moreover, cone specific markers associated with predominantly cone diseases (achromatopsia, cone/cone-rod dystrophy, macular dystrophy) were downregulated, suggesting cones are preferentially affected in *RDH12*-AD models. In addition, 56 genes associated with ocular size, corneal defects and cataracts were dysregulated with 19 downregulated. Further detailed clinical investigation of *RDH12* AD patients may reveal these extra-retinal associations.

Mature retinal organoids resemble more peripheral retina than macula (Cowan et al., 2020), with a well delineated outer limiting membrane and photoreceptor segments exhibiting inner and outer segment characteristics, mitochondria and discs (Zhong et al., 2014). Ultrastructural analysis of mutant retinal organoids revealed shorter photoreceptor segments and reduced cellular numbers than WT controls, consistent with rod-cone dystrophies associated with decreased cone density and shorter cone inner segments as observed in patients by AOSLO (Fingert et al., 2008; Muthiah et al., 2022; Sarkar et al., 2020). While cone defects are observed later in the disease, it cannot be excluded at earlier stages when the patient is potentially pre-symptomatic. Moreover, transmission electronic microscopy of retinal organoids cannot clarify which type of photoreceptors, rods or cones, are impacted. Shorter segments are a sign of photoreceptor degeneration, but no change in ultrastructure of organelles was observed and immunostaining at different timepoints failed to detect a significant increase of cell death in *RDH12*-AD organoids. Moreover, at week 44, two pro-apoptotic genes *MOAP1* and *AIFM2* were downregulated and upregulated, respectively, in *RDH12*-AD retinal organoids (Su et al., 2022; Wu et al., 2002). The equilibrium of these genes might be altered in some cells and lead to their death and thus reduce the number of photoreceptors. Photoreceptor cell death may have occurred between week 18-38, timepoints not examined in this study, hence further ultrastructure analyses at different timepoints during photoreceptor maturation would be needed to assess for longitudinal changes.

*RDH12*-AD retinal organoids are derived from patient cells, heterozygous for c.759del p.(Phe254Leufs^*^24) in *RDH12* (Sarkar et al., 2021, 2020). This frameshift variant creates a premature termination codon (PTC) located 19 nucleotides upstream of the last exon-exon junction in exon 8, predicted to be resistant to nonsense mediated decay (NMD) (Kurosaki et al., 2019).

Transcriptomic analysis reveals *RDH12* expression is decreased by 45% in *RDH12*-AD organoids compared to WT. The presence of mutated *RDH12* mRNA in mutant retinal organoids is suggestive of a partial degradation. The mutated *RDH12* mRNA, which is insensitive to NMD, produces a 40-amino-acid shorter protein, with a 17 amino-acid sequence shared with the two other variants previously reported in autosomal dominant *RDH12*-retinal diseases (Fingert et al., 2008; Muthiah et al., 2022). The WT *RDH12* allele localised to the photoreceptor inner segments (Kurth et al., 2007), including in *RDH12*-AD retinal organoids, was noted at week 18 and week 44. RDH12 was suggested to function as a dimer, based on the orthologue photoreceptor retinal dehydrogenase in *Drosophila melanogaster* (Sarkar and Moosajee, 2019). The alternative C-terminus may function as a dominant-negative protein, sequestering WT protein to reduce its activity.

*RDH12*-AD related RP is characterized by a late-onset milder phenotype as observed in patients affected with *PDE6B*-RP (Kim et al., 2020; Sarkar et al., 2020; Sarkar and Moosajee, 2019). *RDH12* and *PDE6B* are both involved in the vision signal transduction pathway. *PDE6B* expression is unchanged in mature *RDH12*-AD organoids, while *RDH12* is upregulated in *PDE6B* mutant organoids with a missense variant leading to gene upregulation (Gao et al., 2020). This suggests that *PDE6B* regulates *RDH12* expression, but *RDH12* expression does not influence *PDE6B*. Phototransduction in both *PDE6B* and *RDH12*-AD retinal organoid models reveal down-regulation of *ARR3, CNGA3, GNAT2, OPN1LW, PDE6C, PDE6H, RGS9BP* and *RPGRIP1* (Gao et al., 2020).

Further analysis comparing transcriptomic signatures in both RP models could be interesting to investigate common pathways. Transcriptomic analyses at week-44 *RDH12*-AD retinal organoids reveal 79 inherited eye disease-associated genes are differentially expressed, with 48 genes downregulated including 49 associated with IRDs. None of these genes have previously been modelled with patient derived iPSC to study their effect on the retina, but transcriptomic analysis of such retinal organoids may highlight common pathways that could be targeted for more agnostic therapeutic approaches. Of note, *RDH12* expression is restricted to the retina, especially cone and rod photoreceptors (Cowan et al., 2020) but the visual cycle occurs in the retina and the RPE (Kiser et al., 2014). Amongst RPE proteins involved in visual cycle, LRAT converts all-*trans*-retinol into all-*trans*-retinaldehyde, which is transformed into 11-*cis*-retinol by RPE65. RDH5 catalyzes 11-*cis*-retinal biosynthesis from 11-*cis*-retinol, before retinoid transport back in photoreceptors to start a new phototransduction cycle. Mutations in *LRAT* or *RPE65* are associated with Leber congenital amaurosis and variants in *RDH5* lead to *fundus albipunctatus*. However, presently differentiation and culture protocols do not effectively co-culture retinal organoids with RPE (Fathi et al., 2021), limiting the complete identification of affected pathways in AD *RDH12-*RP.

In summary, we have developed the first *RDH12-*AD retinal organoid model and highlight photoreceptor dysfunction and vitamin A pathway defects. The genetic background is highly important in determining disease pathogenicity and severity, hence, isogenic controls would help validate these results (Riordan and Nadeau, 2017). Only four patients have been reported in the literature with AD *RDH12-*RP (Fingert et al., 2008; Muthiah et al., 2022; Sarkar et al., 2020); one unrelated family sharing the same variant studied here (Sarkar et al., 2020), and two other unrelated families, who were heterozygous for c.776del p.(Glu260Argfs*18) or c.763del p.(Val255Serfs*23) variant in *RDH12*, respectively (Fingert et al., 2008; Muthiah et al., 2022). Retinal organoid models from each unrelated family or introduction of the pathogenic variant on the same WT background using CRISPR-Cas9 gene editing would enable further characterisation, electron microscopy and transcriptomic analyses to advance our knowledge of the disease-mechanisms associated with *RDH12-*RP.

## 6 Author Contributions

CM: Conceptualization, Investigation, Methodology, Visualization, Writing – original draft, Writing – review & editing; YZ: Investigation, Visualization, Writing – review & editing; NO: Data curation, Writing – review & editing; PSB: Data curation, Visualization, Writing – review & editing; RC: Formal analysis, Validation, Writing – review & editing; NN: Investigation, Writing – review & editing; HS: Methodology, Writing – review & editing; LT: Methodology, Writing – review & editing; MM: Conceptualization, Funding acquisition, Project administration, Supervision, Writing – original draft, Writing – review & editing.

## 8 Acknowledgments

The authors thank RDH12 patients and their families. They gratefully acknowledge National Institute for Health and Care Research (NIHR) Moorfields BRC, Dhani Tracey-White and Matt Hayes (UCL Institute of Ophthalmology, London) for samples preparation for transmission electron microscopy (TEM), Lucy Collison and the Electron Microscopy Science technology platform (STP) (the Francis Crick Institute, London) for the TEM imaging, Tom Burgoyne for his retinal TEM expertise (UCL Institute of Ophthalmology), Camille Charoy and the Light Microscopy STP (The Francis Crick Institute) and the Bioinformatics and Biostatistics STP (The Francis Crick Institute) for their support and assistance in this work.

## 12 Data Availability Statement

The RNA-seq data generated by this study have been deposited in the NCBI Gene Expression Omnibus (https://www.ncbi.nlm.nih.gov/geo/) under the access code GSE271751, including unprocessed FASTQ files and associated gene count matrices.

