## Supplementary figures and images for "Dominant *RDH12*-retinitis pigmentosa impairs photoreceptor development and cone function in retinal organoids"

### Figure S1

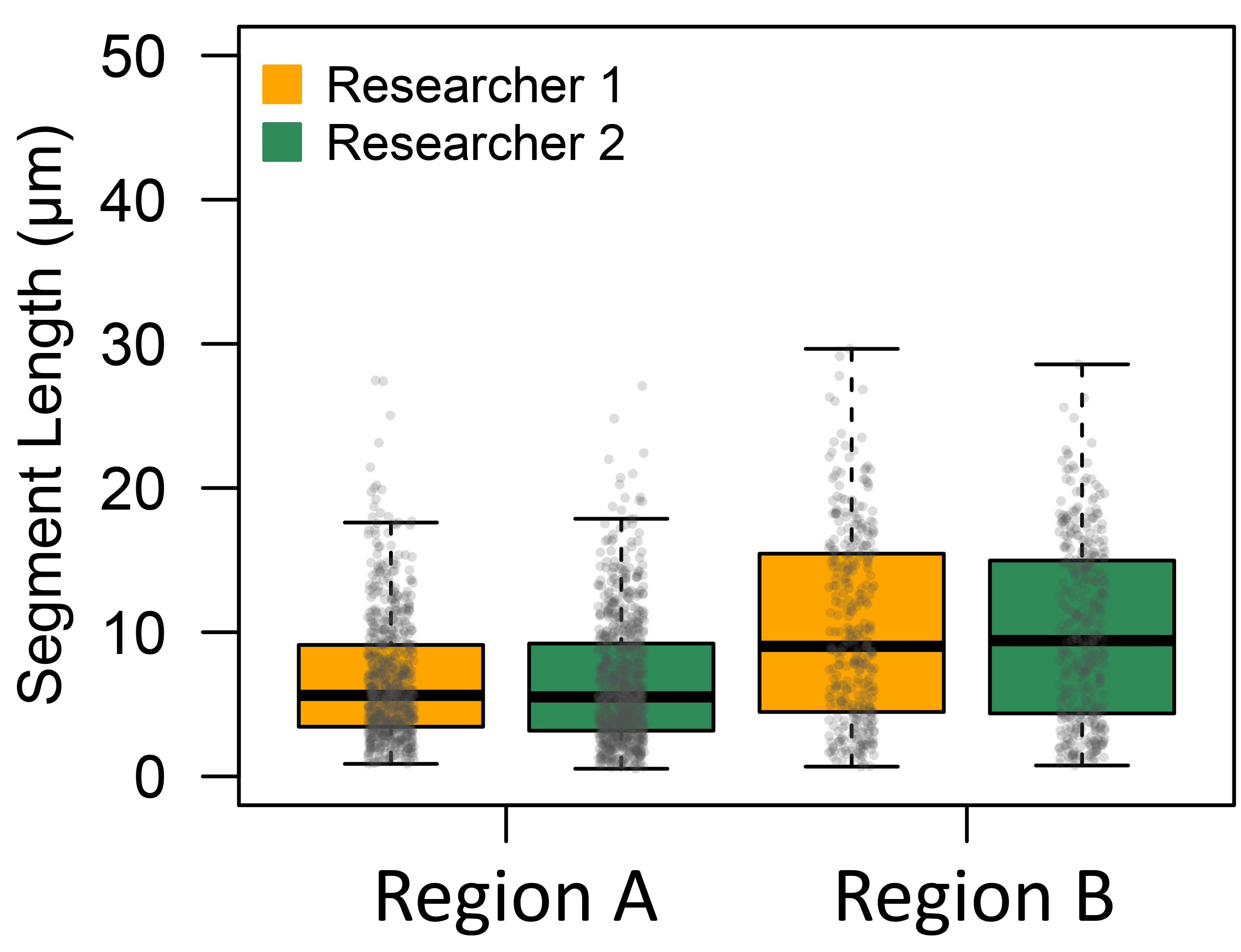
